# buzzdetect: an open-source deep learning tool for automated bioacoustic pollinator monitoring

**DOI:** 10.1101/2025.06.13.659554

**Authors:** Luke E. Hearon, Lillian H.P. Johnson, James Underwood, Chia-Hua Lin, Reed M. Johnson

## Abstract

Ecological studies of pollinators often require long-term and extensive monitoring, posing a significant cost and limitation to research. Traditional sampling methods of observation such as sweep netting and pan trapping provide valuable information on pollinator diversity, but scale poorly when conducting large sampling efforts across space and time. We introduce “buzzdetect”, our tool to apply deep learning models to audio data for conducting passive acoustic monitoring on pollinators, and test our accompanying audio classification model. The model is capable of distinguishing the buzzing of insect flight from environmental noise with a sensitivity of 27% at a precision (true positive rate) of 95%. Application of buzzdetect to recordings made in pumpkin, watermelon, mustard, and soybean reveals differences in timing and intensity of foraging: foraging occurred the earliest in pumpkins and the latest for soybean while foraging activity was most intensive in mustard.

## Introduction

Sampling effort is one of the largest costs of carrying out pollinator research. Ecological systems often have low signal to noise ratios, which demands extensive sampling to overcome. This issue is compounded for studies that seek to characterize patterns in space (e.g., edge effects, habitat fragmentation, population distribution) or in time (e.g., foraging patterns, population dynamics). Studying these effects requires sampling across many points in space or over longer periods of time. The labor and expertise required for sampling can quickly become the limiting factor of a study’s sample size, hampering the ability to discern patterns of pollinator activity with sufficient resolution.

Insect sampling methods can be generally divided into two categories: active and passive. In active methods, the researcher must be present during the course of sampling. Two of the most common active methods of pollinator monitoring are visual observation and sweep netting. Visual observation is appealing for the ability to confirm foraging events on flowers, ensuring that sampled insects are indeed pollinators of the plant in question and not incidental visitors. In contrast, sweep netting affords finer taxonomic resolution and has repeatedly been shown to capture a high abundance and richness of pollinators (Baum and Wallen 2011, Popic et al. 2013, Prendergast et al. 2020). The downside to both of these methods is their time-intensiveness. Because a researcher must be present for the entire duration of sampling, active methods are prohibitively costly for studies that are large in spatial or temporal scale. Passive methods capture insects without the attendance of the researcher. The most common passive method for pollinator monitoring is pan trapping, which allows a single researcher to deploy numerous bee bowls, leave them for the desired duration, and then retrieve and identify samples at leisure. This facilitates extensive sampling across points in space and can capture species across a duration of interest, but it does not improve temporal resolution. To distinguish patterns across time, the researcher must reset the traps at each time point. For example, to measure foraging on an hour-by-hour basis, the researcher must return to every study site at every hour to reset traps, which diminishes the advantage of passive monitoring and may be impossible for large scale studies. Further, the capture of pan traps is itself a function of local floral abundance. When there are few flowers in the local environment, the bowls, which imitate the color of flowers, become relatively more attractive than when surrounding resources are plentiful (Wilson et al. 2008, Baum and Wallen 2011, Popic et al. 2013, Kuhlman et al. 2021, Westerberg et al. 2021). This is a consequential flaw if floral abundance varies across study sites or as the result of an intervention. This bias can be so strong as to completely invert the measured trend of pollinator abundance. St. Clair et al. (2020) found that the number of honey bees captured by pan traps in soybean fields was low during soybean bloom, then increased dramatically when flowering ended. This pattern is implausible, as the authors note, and indeed the opposite trend has been found when sampling with methods not suffering this bias (Souza et al. 2023, Forrester et al. 2024). Finally, after the bouts of sampling have concluded, both sweep netting and pan trapping generate samples of insects that require taxonomic expertise and a significant investment of time to process and identify.

Bioacoustic monitoring offers a promising solution to the above shortcomings. This method detects and identifies organisms by the sounds they produce, allowing recording devices to be deployed at scale as easily as pan traps while offering fine temporal resolution. Because recorders should not attract pollinators, their sampling is not confounded with resource abundance in the local landscape. Most importantly for large-scale studies, the analysis of audio data can be automated through the use of machine learning by building models that identify the sound signatures of species of interest. This is the basis of passive acoustic monitoring, which seeks to answer ecological questions through automated monitoring (Ross et al. 2023). Acoustic monitoring has seen rapid adoption in avian systems (Xie et al. 2023), aquatic systems (Parsons et al. 2022), and whole-soundscape analysis (Nieto-Mora et al. 2023, Turlington et al. 2024). A recent landmark in the field was the release of the Cornell Lab of Ornithology’s powerful avian identification model “BirdNET” (Kahl et al. 2021), which has received hundreds of citations in the few years since its publication—in no small part because of their open-sourcing of their model weights and accessible analysis software (BirdNET Team 2025).

Entomological bioacoustics has seen little development relative to these systems. Kohlberg et al. (2024) recently reviewed publications using bioacoustics for insect detection and identification. The authors identified 49 papers employing automated bioacoustics to monitor insect populations, the earliest being a floppy-disk-based system in 1978, but the majority (28 papers) being published since 2020. Gradišek et al. (2017), Ribeiro et al. (2021) and Ferreira et al. (2023)r tested a number of different algorithms capable of classifying recordings of bumble bee flight or sonication buzzes according to species. However, while these models are impressive, they are not suitable for passive acoustic monitoring because they are not designed to distinguish insect sounds from environmental noise. Rather, they are designed to be run on input audio known to contain a bumble bee buzz. Additionally, to our knowledge, only the models from Ferreira et al. (2023) are publicly available (Ferreira 2025). There also exist some models suitable for passive monitoring. Folliot et al. (2022) produced a convolutional neural network capable of distinguishing insect buzzes from environmental noise, but to the best of our knowledge the model was not made public. Miller-Struttmann et al. (2017) produced a model based on Computational Auditory Scene Analysis (Heise et al. 2017). The model is capable of distinguishing bumble bees from environmental noise, but while the model definition is given in Heise et al. (2017), the model itself is not available for download. The startup company BeeHero produces the Pollinator Insight Platform that appears to operate on a very similar principle as buzzdetect, but it is offered as a commercial service for agriculture and is not developed as a research tool. Of the papers reviewed by Kohlberg et al. (2024), none provided both (i) a readily available algorithm capable of distinguishing insect buzzes from environmental noise and (ii) a tool capable of applying such an algorithm to long-duration field recordings. The goal of this work is to produce a machine learning model capable of detecting insect buzzes for pollinator monitoring, develop an accompanying tool to apply the model to large audio datasets, and apply it to pollinator monitoring in four bee-attractive crops: pumpkin, watermelon, mustard, and soybean.

## Methods

### Training dataset

We produced the training audio for the machine learning model according to bioacoustics methods developed by Forrester et al. (2024). In brief, handheld Sony ICD-PX370 recorders were fastened to plastic electric fence posts, set at the height of the flowers of interest, and left to record for several days (Figure 1). All training audio was produced in soybean fields during flowering in central Ohio. Audio events were manually annotated to serve as training data for the machine learning model (see Table 1 for audio event labels). Audio events were identified by listening to the recordings while inspecting the spectrogram and comparing putative buzzes to reference audio. In total, roughly 24,000 seconds of annotations were generated.

**Figure 1.**
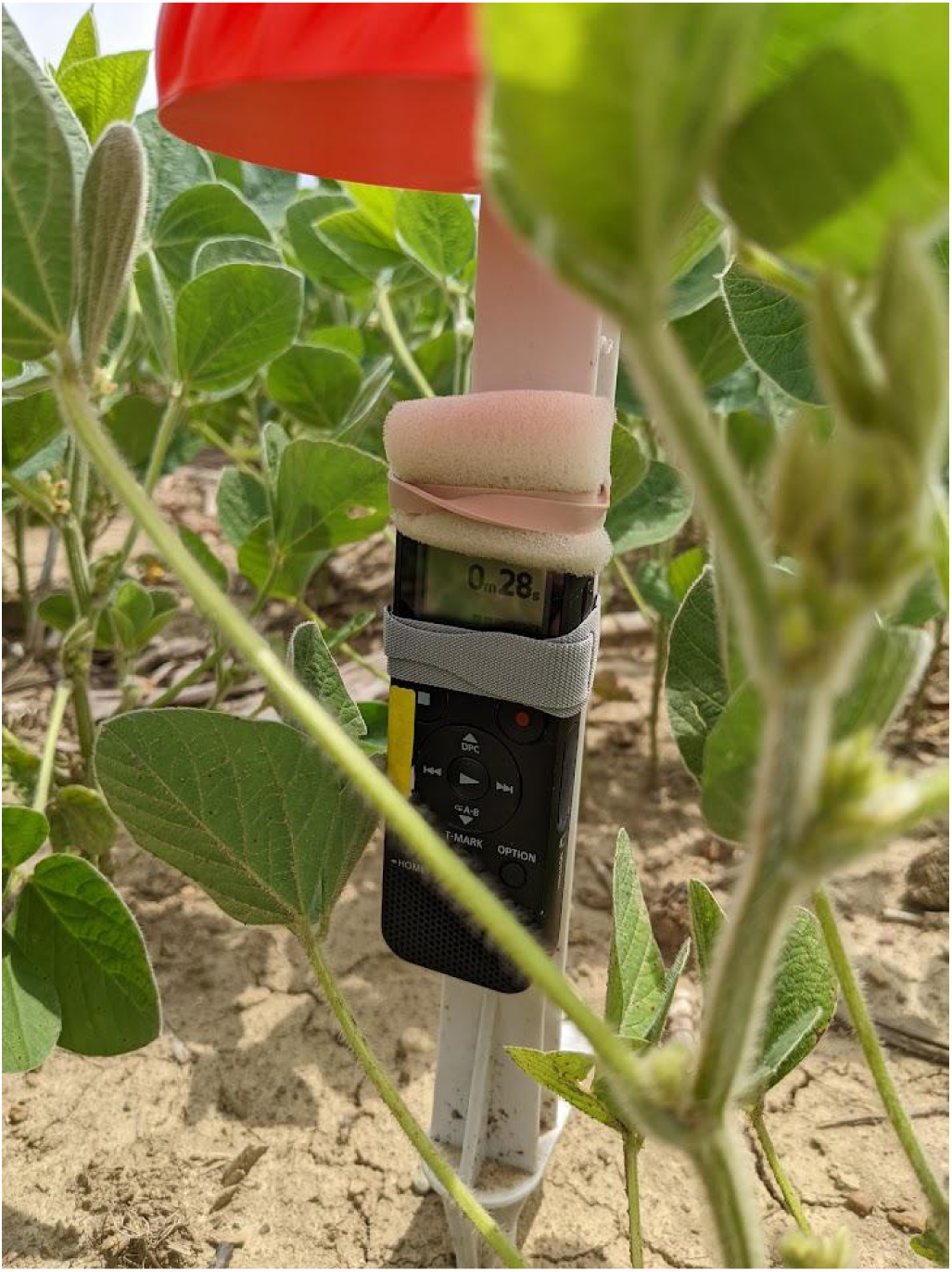
An example of a recorder deployed in soybean. **Alt text:** a Sony recorder strapped to a plastic fence post and deployed low in the foliage of a soybean field.

**Figure 2.**
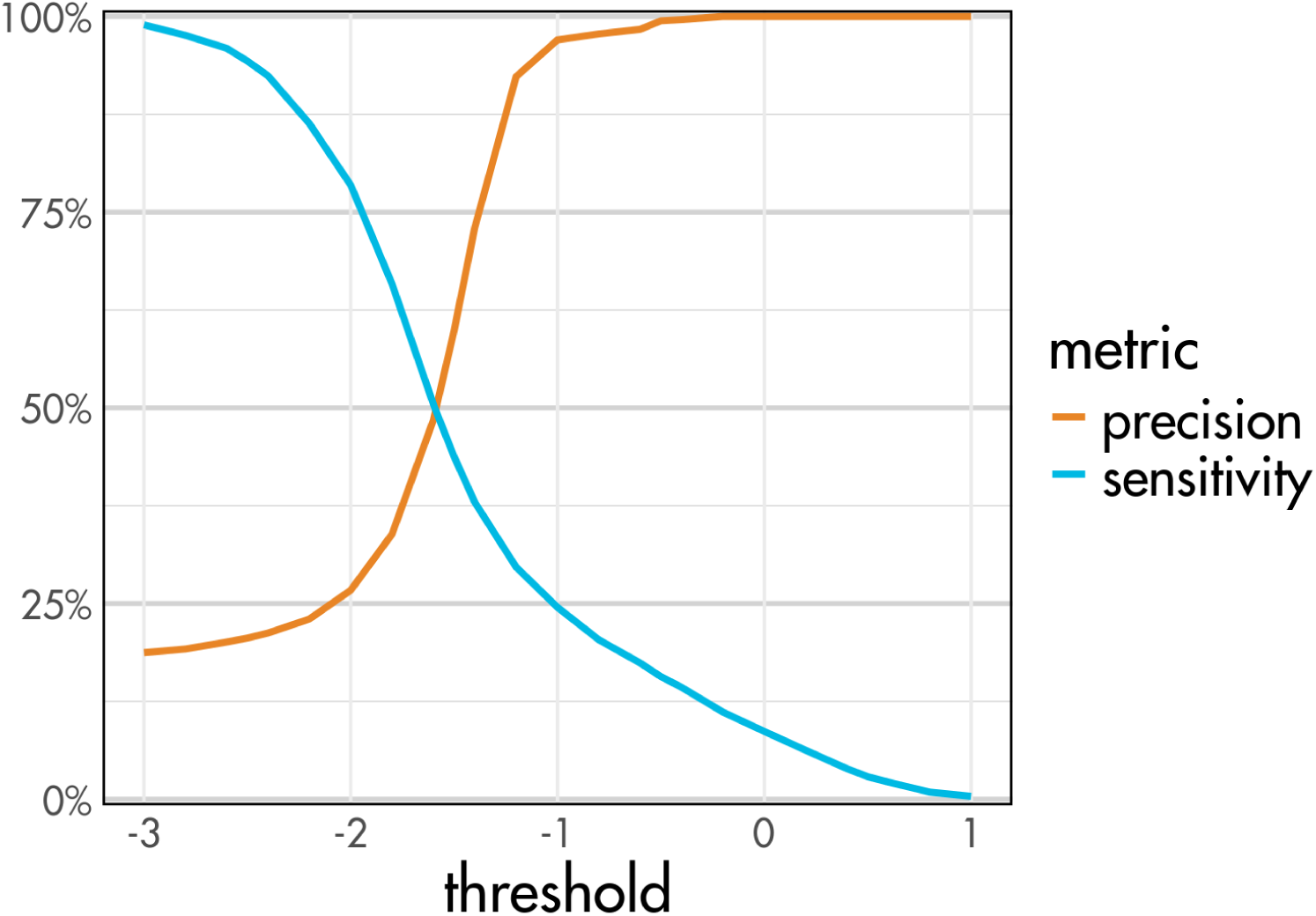
Comparison of model sensitivity (probability of buzz detection given a true buzz in the audio) and precision (probability of a true buzz in the audio given a buzz detection) across a range of threshold values. **Alt text:** A line graph showing that as threshold increases from negative three to positive one, sensitivity decreases from nearly one hundred percent to zero percent and precision increases from twenty percent to one-hundred percent. At a threshold of minus one, the precision is about ninety-five percent and the sensitivity is about twenty-five percent.

**Table 1.** Breakdown of training class volumes. Values in the “label” column correspond to the labels used in buzzdetect model training and output.

| label | description | events | seconds |
| --- | --- | --- | --- |
| ambient_background | Uneventful background noise | 1,503 | 5,322 |
| mech_auto | Cars, trucks, motorcycles | 1,268 | 5,106 |
| mech_plane | Jet planes, propeller planes | 149 | 4,465 |
| ins_buzz | Any insect flight buzz | 1,570 | 2,508 |
| ins_trill | Crickets and cicadas | 336 | 1,915 |
| ambient_noise | Loud ambient noises; scraping foliage, rustling leaves | 816 | 1,414 |
| mech_hum | Monotonous mechanical droning | 68 | 1,283 |
| ambient_rain | Rain and thunder | 54 | 1,141 |
| human | Human vocalization; talking, laughter | 243 | 675 |
| frog | Any calling frog | 201 | 310 |
| mech_siren | Emergency vehicle sirens | 161 | 176 |
| mech_train | Train horns | 83 | 100 |
| bird_goose | Canada goose calls | 27 | 31 |

We labeled around 2,500 seconds worth of buzzing insect flight in the training data. Because it was not possible to rigorously identify insects from the audio alone, we grouped buzzes into three broad categories according to the pitch relative to the buzz of honey bee flight, about 230Hz (Altshuler et al., 2005), for which we had the best reference audio. 8% of the training data for insect buzzes was from buzzes with a pitch higher than honey bees, 58% from medium-pitched buzzes (those that we found to be a good match to our reference audio), and 33% from buzzes with a lower pitch. For training, we conflated all buzz pitches into the single label “ins_buzz” to maximize training data and because we found that classification performance was superior when combining these classes.

### Model training

We employed transfer learning to train a convolutional neural network using a relatively small volume of data. In brief, this technique entails acquiring a pre-trained, general-purpose model and retraining a small part for a specific task rather than attempting to train an entire model from scratch. The general structure for model training followed the “Transfer learning with YAMNet” tutorial by the Google TensorFlow team (TensorFlow Authors 2022). We used the pre-trained audio classifier YAMNet (Plakal and Ellis 2025), which was trained on the AudioSet corpus (Gemmeke et al. 2017) to classify audio into 521 events. However, these events are largely irrelevant to pollinator study systems. We created our own model in order to fine-tune detection for field audio. Our model essentially replaces the final layer of YAMNet, which performs classification. We use the values from the second-to-last layer of YAMNet (the embedding layer, see TensorFlow Authors 2022) as the input and connect these values to 8 output neurons corresponding to our events of interest (see Table 1). We then trained our model by applying YAMNet to our training dataset, extracting values from the second-to-last layer, and training our model on the extracted values.

### buzzdetect

Accompanying the model, we developed a Python-based tool to facilitate the analysis of large volumes of audio data. We call the tool “buzzdetect” (stylized in the lower case). To accelerate analysis, buzzdetect employs parallel processing and optional processing on GPU. Audio files to be analyzed should not require any preprocessing: we use the packages soundfile (Bechtold 2025) to read a wide variety of file formats and librosa (McFee et al. 2023) to resample audio in-memory to the appropriate sample rate. On budget hardware (Intel Core i7-2600 8-core CPU, NVIDIA GeForce GTX 1650 GPU, 7200 RPM Western Digital HDD), we see an analysis rate of roughly 1,400x, analyzing an hour of audio in under three seconds.

YAMNet segments input audio into discrete 0.96-second “frames” and performs classification on each frame. Thus, for each input audio file, buzzdetect outputs model results to a CSV file as follows: each row corresponds to a single audio frame, a “start” column records the timestamp of the start of the frame in seconds, and a column for each output neuron records the activation value of the neuron for that audio frame.

An output neuron’s activation is a measure of the model’s confidence that the neuron’s corresponding event is present in the input data. Thus, where the ‘ins_buzz’ neuron activation is high, we expect to hear the buzz of insect flight. The researcher then sets a threshold above which to call detections. The choice of a threshold value is a subjective one; higher or lower thresholds may be desired depending on how the researcher values the trade-off between the sensitivity and its precision. We use sensitivity to mean the probability of a detection given the occurrence of insect flight buzzing in the frame and precision to mean the probability that a buzz is present in the frame given a detection (the true positive rate). A higher threshold is more conservative; it will produce a higher precision (called detections are more likely to be true) at the cost of sensitivity (true buzzes are less likely to be called).

### Model validation

To evaluate the model’s performance in a realistic, long-term context, we selected a small subset of audio files from our previous unpublished work in four crops planted in central Ohio: pumpkin, watermelon, mustard, and soybean. None of the audio used for evaluation was part of the model’s training set and all of the validation recordings were conducted at different sites than the training recordings. This provides a truly out-of-sample test set with real-world audio event distributions and distinct environmental noise profiles.

Each of these studies followed the bioacoustic methods developed by Forrester et al. (2024) and described in the “Training dataset” section above. From each experiment, we selected audio from four randomly selected recorders on a single day near the middle of the flowering period of the crop: August 8, 2024 for pumpkin, July 27, 2024 for watermelon, September 9, 2024 for mustard, and August 10, 2023 for soybean. This produced four 24-hour recordings from each crop for a total of 16 recordings. We then selected one recording per crop and manually annotated the audio events contained in the first five minutes of every hour of the recording from 6:00am to 8:00pm. This produced a total of 60 manually annotated time periods for comparison to the buzzdetect model. Our annotations for the validation data followed the same labeling scheme as for the training data. We matched every frame of the buzzdetect results to their corresponding annotations in order to calculate sensitivity and precision. In this way, we evaluated overall model performance as well as performance for each crop and for each buzz pitch.

### Application

To simulate a research application of buzzdetect, we applied the model to the full 24-hour recordings. We used a threshold corresponding to 95% specificity, as determined by our validation results, and calculated the detection rate across time by binning results into 10-minute bins and dividing the number of positive frames by the total number of frames in the bin. Using the R (R Core Team 2025) package ggplot2 (Wickham 2016), we plotted the trend of each recorder across the day to show the pattern of diel foraging in each crop.

We tested for differences in the average foraging intensity in each crop by calculating the average detection rate across the course of the day and regressing this against crop type in a beta regression using the R package betareg (Cribari-Neto and Zeileis 2010). To correct for differences in the model’s sensitivity in different crops, we scaled each recorder’s detection rate according to crop sensitivity as determined during model validation. Differences were tested in a pairwise post-hoc test using the pairs.emmGrid function of the package emmeans (Lenth 2024).

We tested for differences in the timing of foraging in each crop by determining the 10-minute bin with peak foraging for each recorder. We expressed the time of day as a decimal with 0 and 1 representing midnight and 0.50 representing noon. Time of peak foraging was regressed against crop type in a linear model using the base R function lm (R Core Team 2025) and differences between crops were again tested with a pairwise post-hoc test using the pairs.emmGrid function of the package emmeans (Lenth 2024).

## Results

### Annotation

We observed a continuous hum of insect buzzing in mustard from 10:00am–2:00pm and in soybean from 12:00pm–2:00pm. The spectrogram for this hum was enriched in bands that matched the medium buzz pitch; all audio was considered positive for medium buzzes at those times.

### Model Validation

Averaged across all crops and buzz pitches, the model could detect buzzes with a 95% precision and 27% sensitivity at a threshold of −1.1. At this threshold, the model was most sensitive to medium-pitched buzzes (26%) and low-pitched buzzes (24%), and insensitive to high-pitch buzzes (0%). This relationship was similar across thresholds.

Model performance varied by crop (Figure 3). At the average 95% precise threshold of −1.1, the precision in pumpkin was 89% and the sensitivity was 9%; in watermelon, the precision was 80% and sensitivity was 12%; in mustard, the precision was 95% and the sensitivity was 31%; in soybean, the precision was 96% and the sensitivity was 26%.

**Figure 3.**
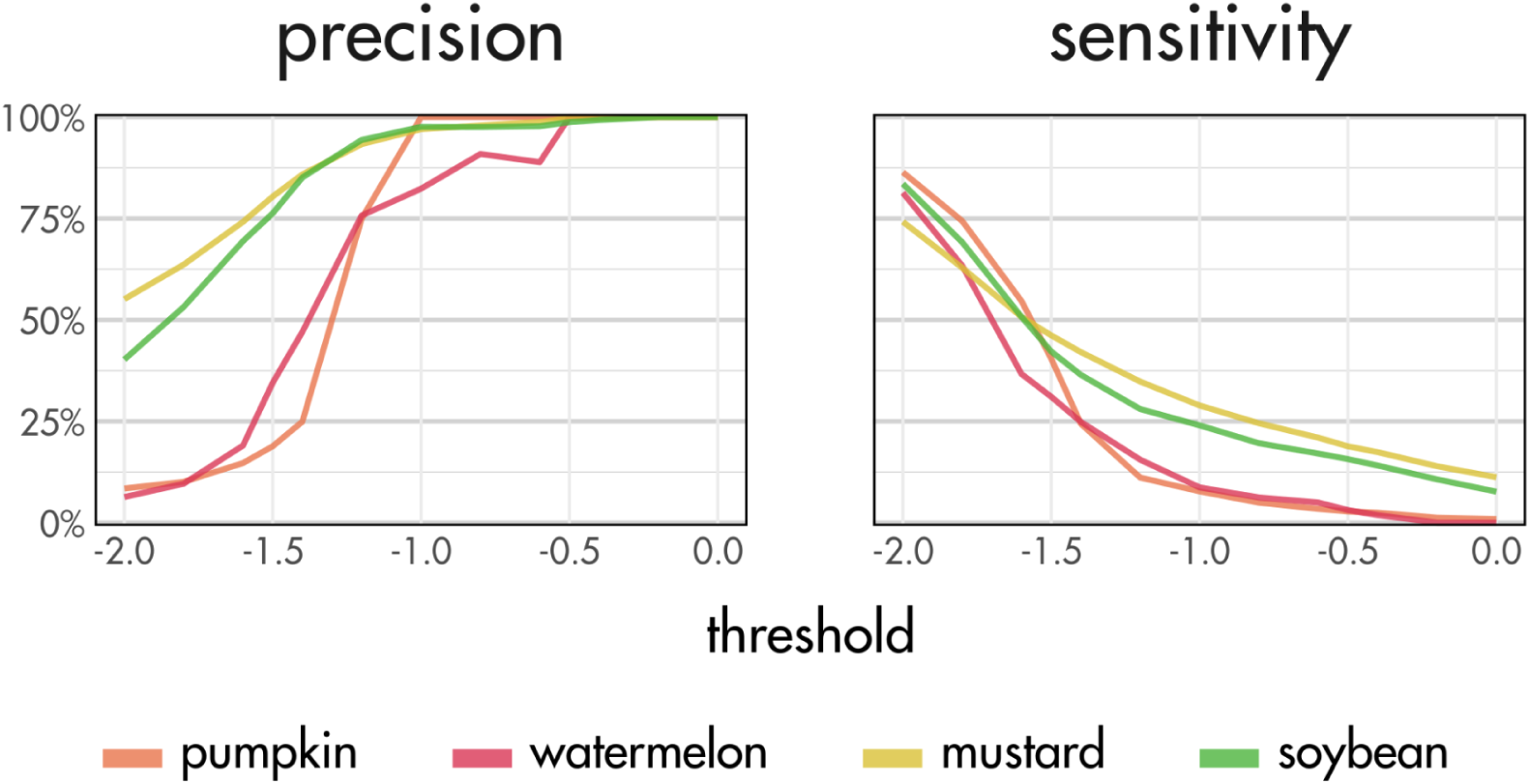
Comparison of sensitivity and precision in different crops. **Alt text:** Two line graphs. The first shows that at a threshold of negative one, precision is above ninety-five percent for all crops but watermelon, which has about eighty percent precision. The second shows that sensitivity is highest for mustard and soybean around twenty five percent and lower for watermelon and pumpkin around ten percent.

### Application

Foraging activity varied considerably between recorders in the same crop (Figure 4). One recorder in watermelon showed much greater activity than the others, with 3,202 detections over the 24-hour period while the other recorders averaged around 340. This difference was genuine, not a model artifact, as a review of the audio around peak foraging confirmed the presence of many buzzes. Even in mustard where the four recorders showed the most consistent trend in foraging activity, detection rates spanned a range of roughly 0.50 during the period of peak activity.

**Figure 4.**
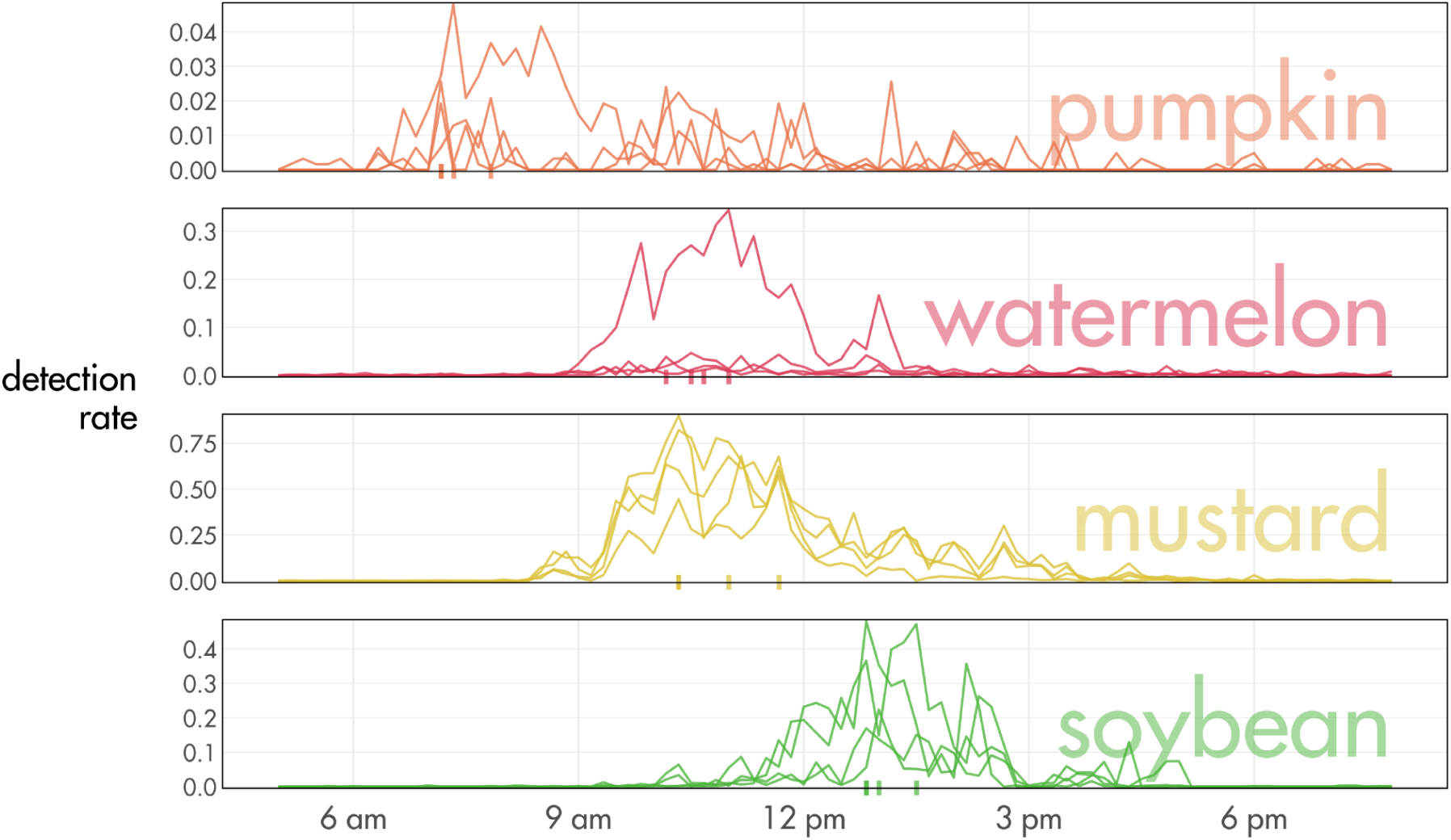
Activity curves for each of the four recorders for each crop. Detection rate is calculated in 10-minute bins. Hash marks on the X axis represent peak foraging time of one of the recorders. **Alt text:** A line graph showing that pumpkin has high activity around eight in the morning, watermelon and mustard have high activity shortly before noon, and soybean has high activity around one in the afternoon.

Mustard showed significantly higher foraging than soybean and pumpkin (both *z* > 5.9, both *p* < 0.001), but not higher foraging than watermelon (*z* = 2.1, *p =* 0.14). The lack of difference with watermelon was driven by one recorder with an unusually high detection rate; after removing the data from this recorder, the difference between mustard and watermelon was significant (*z* = 11.4, *p* < 0.001). The other crops did not significantly differ from one another in average detection rate.

These results also reveal differences in the timing of foraging between these crops. With the exception of watermelon and mustard (*t* = −0.4, *p* = 0.98), each crop differed from all other crops in the timing of peak foraging (all *p* < 0.05).

## Discussion

The diel patterns of foraging in different crops captured by buzzdetect are corroborated by existing literature. Foraging in watermelon (Njoroge et al. 2010, Di Trani et al. 2022, Korichi et al. 2025), pumpkin (Ali et al. 2014, Phillips and Gardiner 2015) and other cucurbits has been shown to peak early in the morning and taper off by midday, while foraging in soybean sees a more gradual onset and offset later in the day (Chiari et al. 2005, Blettler et al. 2018, Jung et al. 2020, Souza et al. 2023, Forrester et al. 2024). The trend in mustard is less distinct, but a midday bloom with high visitation is also supported by existing literature (Mandal et al. 2018, Gautam et al. 2022, Adlin Prajula et al. 2023, Priyadarshini et al. 2025). In part, these differences are likely due to floral attractiveness and phenology, but these data were collected in different locations at different times of the year. With this small sample, we seek to demonstrate the potential of buzzdetect to identify foraging patterns rather than to draw firm conclusions as to their drivers in these particular crops.

An important caveat in interpreting the sensitivity value is that it is calculated in terms of audio frames, not buzzes. However, a buzz might span multiple frames. Insect buzzes in our annotations lasted about 1.5 seconds on average, which would be expected to span 2.6 frames given a random position in the audio framing. Thus, using our model’s per-frame sensitivity of 27% the probability of missing a detection in all three frames is closer to 0.73^2.6^ = 0.45, a per-buzz sensitivity of 55%.

We found that the general trend of activity over the course of the day looks similar even at remarkably low sensitivities (Figure 5). In part, this is likely due to the difference between per-frame and per-buzz sensitivity. Additionally, over the course of an entire day, even a small sample of buzzes is sufficient to capture an informative trend. At a precision of 95%, the number of detections our model produced in a day averaged across the four crops was about 2,800. Comparing this to the normal abundance of capture in pan traps and sweep nets, the realized sensitivity of this method may be comparable to other sampling methods. For example, Hudson et al. (2020) found that only about 20% of bees that approached a pan trap were successfully captured by the trap.

**Figure 5.**
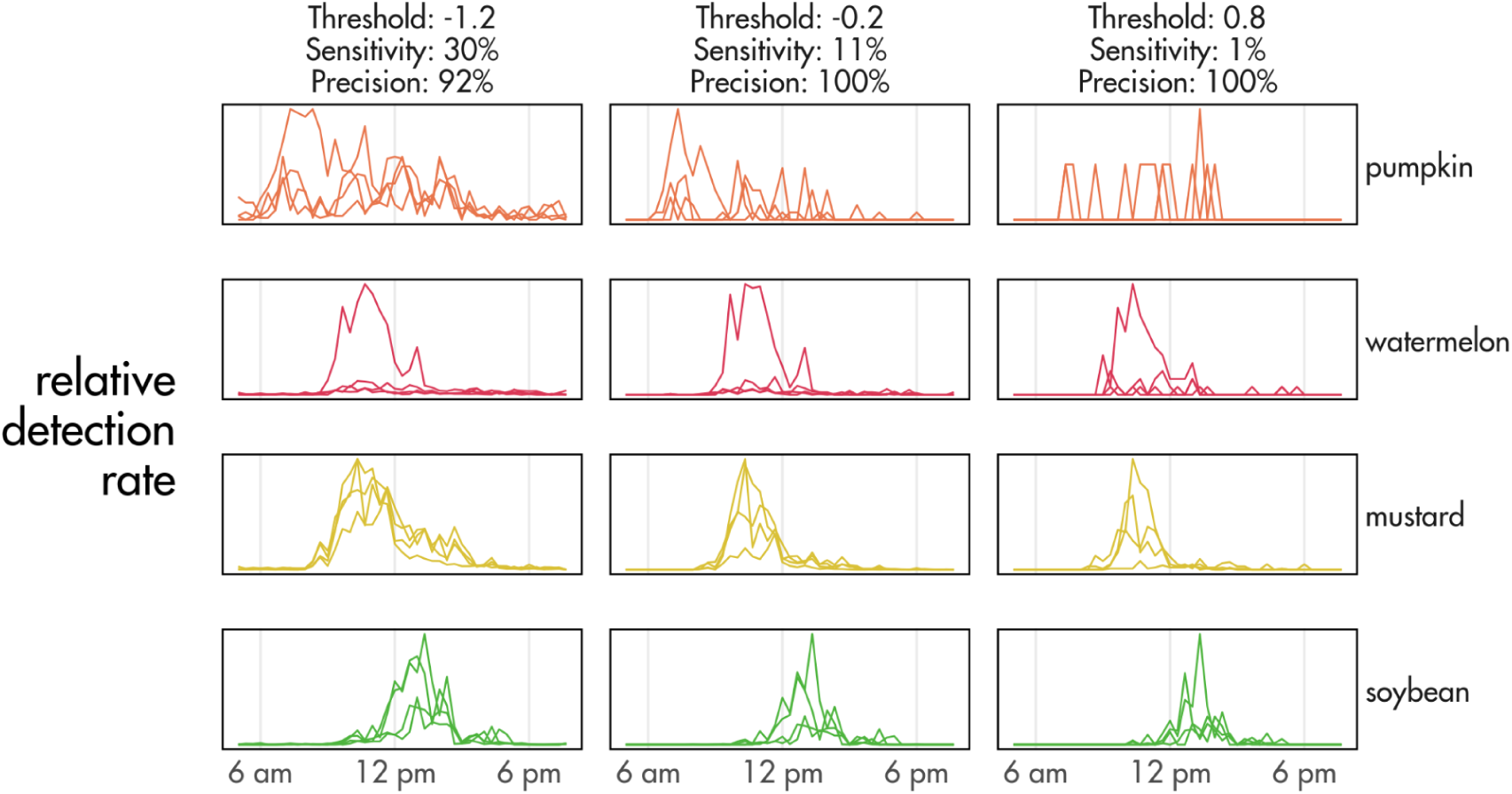
Trends in activity curves are similar even when sensitivity drops very low. The Y-axis is lacking values because detection rates have relativized for each panel to better visualize the trend. **Alt text:** A line graph showing the same trends as in Figure 4, now repeated multiple times at different threshold values. As the threshold increases, sensitivity decreases to one percent. However, the foraging trends look similar even as sensitivity decreases.

### Considerations

While our model can detect insect buzzes, it is not able to identify the insect producing the buzz. Distinguishing between species by their flight buzzes is a more challenging task than discriminating buzzes from environmental noise, and preliminary testing showed that our volume of training data was not sufficient to train a model to make such a distinction. Prior work has produced models capable of classifying insects given a positive example of a buzz (Gradišek et al. 2017, Ribeiro et al. 2021, Ferreira et al. 2023), and buzzdetect could serve as a first-pass to extract portions of audio positive for buzzes to then be identified by these more specific models.

While the model’s sensitivity to low and medium buzzes was comparable, high buzzes were very unlikely to produce a detection. This implies that model performance will differ between pollinator assemblages. Indeed, sensitivity varied by crop type (Figure 3), which appears to be driven by differences in pollinator composition (Table 2). The model showed least sensitivity in the crops with the most high-pitched buzzes. While it may be possible to address this shortcoming by further sampling of underrepresented classes, a similar problem is inherent to all bioacoustics applications: louder and more active organisms are inherently easier to detect. For a given level of sound energy, the buzz of a bumble bee propagates farther than the buzz of a mosquito. The bird that sings incessantly will produce more detections than the one that remains silent.

**Table 2.** The distribution of buzz pitches from manual annotation, given in seconds and as a percentage of the total buzzes for that crop.

| crop | buzz pitch |  |  | total (sec) |
| --- | --- | --- | --- | --- |
|  | high (sec) | medium (sec) | low (sec) |  |
| pumpkin | 88 (42.5%) | 67 (32.3%) | 53 (25.3%) | 208 |
| watermelon | 12 (11.1%) | 79 (75.5%) | 14 (13.3%) | 105 |
| mustard | 9 (0.4%) | 2138 (99.6%) | 0 | 2146 |
| soybean | 1 (0.1%) | 1138 (99.9%) | 0 | 1139 |

We observed a continuous hum of insect buzzing in mustard from 10:00am–2:00pm and in soybean from 12:00pm–2:00pm. The spectrogram for this hum was enriched in bands that matched the medium buzz pitch; all audio was considered positive for medium buzzes at those times.

Imperfect detection pervades all sampling methods (Chen et al. 2013, Kellner and Swihart 2014), but it is not without redress (Royle et al. 2005, Guillera-Arroita 2017). It is beyond the scope of this article to fully address imperfect detection arising from bioacoustics (see Ducrettet et al. 2025 for detailed discussion), but we offer two recommendations for interpreting buzzdetect results. First, a small subsample of audio from an experiment can be manually validated to calibrate model results for differential sensitivity, as we did in our average foraging analysis. Second, bioacoustic monitoring can be supplemented with smaller bouts of traditional sampling. A light sampling effort with visual observation or sweep netting could provide a ground-truth with which to compare acoustic detections, while still capitalizing on the labor savings of automated monitoring. These methods complement one another: bioacoustics provides intensive, time-rich, automatic monitoring while traditional trapping provides rich taxonomic information and direct observation of pollinator activity.

Like all sampling methods, our passive acoustic monitoring method is not without trade-offs. In some applications, these trade-offs may not meaningfully impact the results: deployment in patches where one species is the predominant forager, where pollinator assemblage is constant, or in order to examine the timing, but not intensity, of insect activity. In other applications, we suggest that bioacoustic sampling be supplemented with other sampling methods. Passive acoustic monitoring is a rapidly evolving field and we believe buzzdetect is a promising first step in enabling automated, long-context, and large-scale monitoring of pollinators. Buzzdetect is available at github.com/OSU-Bee-Lab/buzzdetect.

## Data availability statement

All data and code needed to reproduce the results in this manuscript are available at:

Hearon, L., Johnson, L. H. P., Underwood, J., Lin, C.-H., & Johnson, R. M. (2025). Data and code for “buzzdetect: an open-source deep learning tool for automated bioacoustic pollinator monitoring” [Data set]. Zenodo. https://doi.org/10.5281/zenodo.15644084

The buzzdetect source code and current models are available at https://github.com/OSU-Bee-Lab/buzzdetect

## Acknowledgments

This work was funded by the Agriculture and Food Research Initiative of the US Department of Agriculture’s National Institute of Food and Agriculture under grant 2022-67019-36437, Project Apis.m Healthy Hives Research Initiative, The One Hive Foundation under grant 027798, and The Ohio State University College of Food, Agricultural, and Environmental Sciences’s Internal Grant Program (Project #2023-013). Research support was provided by state and federal funds appropriated to the Ohio State University, College of Food, Agricultural, and Environmental Sciences, Ohio Experiment Station (OHO01277; OHO01355-MRF).

We thank Adam Foster for his assistance in designing and executing the study to collect audio in pumpkin and watermelon and Lauren Tarver for her assistance in audio collection. We thank Breh Ruger, Ashley Leach, Laura Lindsey, and Alan Geyer for the support they offered by allowing us access to crop fields for recording.

## Works Cited

Adlin Prajula J, Acharya V, Syed Mohamed Ibrahim S. 2023. Insect Pollinators of Mustard and their Foraging Behaviour. Indian Journal of Entomology.:1–4. 10.55446/IJE.2023.1096

Ali M, Saeed S, Sajjad A, et al. 2014. Exploring the best native pollinators for pumpkin (cucurbita pepo) production in punjab, pakistan. Pakistan Journal of Zoology. 46(2). https://www.proquest.com/docview/1553123108/citation/712AFFBFA6564DD3PQ/1.

Altshuler DL, Dickson WB, Vance JT, et al. 2005. Short-amplitude high-frequency wing strokes determine the aerodynamics of honeybee flight. Proceedings of the National Academy of Sciences. 102(50):18213–18218. 10.1073/pnas.0506590102

Baum KA, Wallen KE. 2011. Potential bias in pan trapping as a function of floral abundance. Journal of the Kansas Entomological Society. 84(2):155–159. 10.2317/JKES100629.1

Bechtold B. 2025. Bastibe/python-soundfile. https://github.com/bastibe/python-soundfile.

BirdNET Team. 2025. BirdNET-Team. https://github.com/birdnet-team/BirdNET-Analyzer.

Blettler DC, Fagúndez GA, Caviglia OP. 2018. Contribution of honeybees to soybean yield. Apidologie. 49(1):101–111. 10.1007/s13592-017-0532-4

Chen G, Kéry M, Plattner M, et al. 2013. Imperfect detection is the rule rather than the exception in plant distribution studies. Journal of Ecology. 101(1):183–191. 10.1111/1365-2745.12021

Chiari WC, Toledo V de AA de, Ruvolo-Takasusuki MCC, et al. 2005. Pollination of soybean (Glycine max L. Merril) by honeybees (Apis mellifera L.). Brazilian Archives of Biology and Technology. 48:31–36. 10.1590/S1516-89132005000100005

Cribari-Neto F, Zeileis A. 2010. Beta regression in R. Journal of Statistical Software. 34(2):1–24. 10.18637/jss.v034.i02

Di Trani JC, Ramírez VM, Añino Y, et al. 2022. Environmental conditions and bee foraging on watermelon crops in Panama. Journal of Animal Behaviour and Biometeorology. 10(4):2234. 10.31893/jabb.22034

Ducrettet M, Linossier J, Sueur J, et al. 2025. Bridging Passive Acoustic Monitoring and Essential Biodiversity Variables with detectability. 10.22541/au.174308491.17489353/v1

Ferreira AIS, da Silva, Nádia Felix Felipe, Mesquita, Fernanda Neiva, et al. 2023. Automatic acoustic recognition of pollinating bee species can be highly improved by Deep Learning models accompanied by pre-training and strong data augmentation. Frontiers in Plant Science. 14. 10.3389/fpls.2023.1081050

Ferreira ALS. 2025. Alefiury/transformers-bee-species-acoustic-recognition. https://github.com/alefiury/Transformers-Bee-Species-Acoustic-Recognition.

Folliot A, Haupert S, Ducrettet M, et al. 2022. Using acoustics and artificial intelligence to monitor pollination by insects and tree use by woodpeckers. Science of The Total Environment. 838. 10.1016/j.scitotenv.2022.155883

Forrester KC, Lin C-H, Johnson RM. 2024. Measuring factors affecting honey bee (hymenoptera: Apidae) attraction to soybeans using bioacoustics monitoring. Journal of Insect Science. 24(2):20. 10.1093/jisesa/ieae036

Gautam RK, Shuyi G, Uniyal VP. 2022. Comparative foraging behaviour and pollination efficiency of *Apis laboriosa* S. and *Apis cerana* F. on black mustard (*Brassica nigra* L.) in Western Himalaya, India. Current Science. 122(7):840. 10.18520/cs/v122/i7/840-845

Gemmeke JF, Ellis DPW, Freedman D, et al. 2017. 2017 IEEE international conference on acoustics, speech and signal processing (ICASSP). p. 776–780. 10.1109/ICASSP.2017.7952261

Gradišek A, Slapničar G, Šorn J, et al. 2017. Predicting species identity of bumblebees through analysis of flight buzzing sounds. Bioacoustics-the International Journal of Animal Sound and Its Recording. 26(1):6376. 10.1080/09524622.2016.1190946

Guillera-Arroita G. 2017. Modelling of species distributions, range dynamics and communities under imperfect detection: advances, challenges and opportunities. Ecography. 40(2):281–295. 10.1111/ecog.02445

Heise D, Miller-Struttmann N, Galen C, et al. 2017. 2017 IEEE sensors applications symposium (SAS). Glassboro, NJ, USA: IEEE. p. 1–5. 10.1109/SAS.2017.7894089

Hudson J, Horn S, Hanula JL. 2020. Assessing the Efficiency of Pan Traps for Collecting Bees (Hymenoptera: Apoidea). Journal of Entomological Science. 55(3):321–328. 10.18474/0749-8004-55.3.321

Jung AH, Perini CR, Valmorbida I, et al. 2020. Foraging, spatial distribution and the effect of honeybees on soybean yield. Australian Journal of Crop Science.(14(12):2020):1983–1990. 10.21475/ajcs.20.14.12.2855

Kahl S, Wood CM, Eibl M, et al. 2021. BirdNET: A deep learning solution for avian diversity monitoring. Ecological Informatics. 61:101236. 10.1016/j.ecoinf.2021.101236

Kellner KF, Swihart RK. 2014. Accounting for Imperfect Detection in Ecology: A Quantitative Review. Buckel J, editor. PLoS ONE. 9(10):e111436. 10.1371/journal.pone.0111436

Kohlberg AB, Myers CR, Figueroa LL. 2024. From buzzes to bytes: A systematic review of automated bioacoustics models used to detect, classify and monitor insects. Journal of Applied Ecology. 61(6). 10.1111/1365-2664.14630

Korichi Y, Karima K-G, Malika A-S. 2025. Diversity and foraging behavior assessment of watermelon pollinators in Tizi-Ouzou area (Algeria). Journal of Apicultural Research. 64(1):159–166. 10.1080/00218839.2023.2250454

Kuhlman MP, Burrows S, Mummey DL, et al. 2021. Relative bee abundance varies by collection method and flowering richness: Implications for understanding patterns in bee community data. Ecological Solutions and Evidence. 2(2):e12071. 10.1002/2688-8319.12071

Lenth RV. 2024. Emmeans: Estimated marginal means, aka least-squares means. https://CRAN.R-project.org/package=emmeans.

Mandal E, Amin MR, Rahman H, et al. 2018. Abundance and foraging behavior of native insect pollinators and their effect on mustard (Brassica juncea L.). Bangladesh Journal of Zoology. 46(2):117–123. 10.3329/bjz.v46i2.39045

McFee B, McVicar M, Faronbi D, et al. 2023. Librosa/librosa: 0.10.0. 10.5281/zenodo.7657336

Miller-Struttmann NE, Heise D, Schul J, et al. 2017. Flight of the bumble bee: Buzzes predict pollination services. PLOS ONE. 12(6):e0179273. 10.1371/journal.pone.0179273

Nieto-Mora DA, Rodríguez-Buritica S, Rodríguez-Marín P, et al. 2023. Systematic review of machine learning methods applied to ecoacoustics and soundscape monitoring. Heliyon. 9(10):e20275. 10.1016/j.heliyon.2023.e20275

Njoroge N, Gemmill B, Bussmann R, et al. 2010. Diversity and efficiency of wild pollinators of watermelon (Citrullus lanatus (Thunb.) Mansf.) at Yatta (Kenya). Journal of Applied Horticulture. 12(01):35–41. 10.37855/jah.2010.v12i01.07

Parsons MJG, Lin T-H, Mooney TA, et al. 2022. Sounding the Call for a Global Library of Underwater Biological Sounds. Frontiers in Ecology and Evolution. 10:810156. 10.3389/fevo.2022.810156

Phillips BW, Gardiner MM. 2015. Use of video surveillance to measure the influences of habitat management and landscape composition on pollinator visitation and pollen deposition in pumpkin ( *Cucurbita pepo*) agroecosystems. PeerJ. 3:e1342. 10.7717/peerj.1342

Plakal M, Ellis D. 2025. Google yamnet Kaggle.

Popic TJ, Davila YC, Wardle GM. 2013. Evaluation of Common Methods for Sampling Invertebrate Pollinator Assemblages: Net Sampling Out-Perform Pan Traps. PLOS ONE. 8(6):e66665. 10.1371/journal.pone.0066665

Prendergast KS, Menz MHM, Dixon KW, et al. 2020. The relative performance of sampling methods for native bees: an empirical test and review of the literature. Ecosphere. 11(5):e03076. 10.1002/ecs2.3076

Priyadarshini P, Satapathy SN, Sahoo BK, et al. 2025. Diversity and Relative Abundance of Insect Pollinators on Mustard, Brassica Juncea L. HEXAPODA.:1–6. 10.55446/hexa.2025.572

R Core Team. 2025. R: A language and environment for statistical computing. Vienna, Austria: R Foundation for Statistical Computing. https://www.R-project.org/.

Ribeiro AP, Silva NFF da, Mesquita FN, et al. 2021. Machine learning approach for automatic recognition of tomato-pollinating bees based on their buzzing-sounds. PLOS Computational Biology. 17(9):e1009426. 10.1371/journal.pcbi.1009426

Ross SRP-J, O’Connell DP, Deichmann JL, et al. 2023. Passive acoustic monitoring provides a fresh perspective on fundamental ecological questions. Functional Ecology. 37(4):959–975. 10.1111/1365-2435.14275

Royle JA, Nichols JD, Kéry M. 2005. Modelling occurrence and abundance of species when detection is imperfect. Oikos. 110(2):353–359. 10.1111/j.0030-1299.2005.13534.x

Souza EP, Degrande PE, Barbosa VO, et al. 2023. Temporal dynamics of Apis mellifera (Hymenoptera: Apidae) during flowering in indeterminate soybean (Glycine max). Anais da Academia Brasileira de Ciências. 95(4):e20191214. 10.1590/0001-3765202320191214

St. Clair AL, Dolezal AG, O’Neal ME, et al. 2020. Pan Traps for Tracking Honey Bee Activity-Density: A Case Study in Soybeans. Insects. 11(6):366. 10.3390/insects11060366

TensorFlow Authors. 2022. Transfer learning with YAMNet for environmental sound classification. https://tensorflow.org/tutorials/audio/transfer_learning_audio

Turlington K, Suárez-Castro AF, Teixeira D, et al. 2024. Exploring the relationship between the soundscape and the environment: A systematic review. Ecological Indicators. 166:112388. 10.1016/j.ecolind.2024.112388

Westerberg L, Berglund H-L, Jonason D, et al. 2021. Color pan traps often catch less when there are more flowers around. Ecology and Evolution. 11(9):3830–3840. 10.1002/ece3.7252

Wickham H. 2016. ggplot2: Elegant graphics for data analysis. Springer-Verlag New York. https://ggplot2.tidyverse.org.

Wilson JS, Griswold T, Messinger OJ. 2008. Sampling Bee Communities (Hymenoptera: Apiformes) in a Desert Landscape: Are Pan Traps Sufficient? Journal of the Kansas Entomological Society. 81(3):288–300. 10.2317/JKES-802.06.1

Xie J, Zhong Y, Zhang J, et al. 2023. A review of automatic recognition technology for bird vocalizations in the deep learning era. Ecological Informatics. 73:101927. 10.1016/j.ecoinf.2022.101927

